# Resources for the comprehensive discovery of functional RNA elements

**DOI:** 10.1101/030486

**Authors:** Balaji Sundararaman, Lijun Zhan, Steven Blue, Rebecca Stanton, Keri Elkins, Sara Olson, Xintao Wei, Eric L. Van Nostrand, Stephanie C. Huelga, Brendan M. Smalec, Xiaofeng Wang, Eurie L. Hong, Jean M. Davidson, Eric Lecuyer, Brenton R. Graveley, Gene W. Yeo

## Abstract

Transcriptome-wide maps of RNA binding protein (RBP)-RNA interactions by immunoprecipitation (IP)-based methods such as RNA IP (RIP) and crosslinking and IP (CLIP) are key starting points for evaluating the molecular roles of the thousands of human RBPs. A significant bottleneck to the application of these methods in diverse cell-lines, tissues and developmental stages, is the availability of validated IP-quality antibodies. Using IP followed by immunoblot assays, we have developed a validated repository of 438 commercially available antibodies that interrogate 365 unique RBPs. In parallel, 362 short-hairpin RNA (shRNA) constructs against 276 unique RBPs were also used to confirm specificity of these antibodies. These antibodies can characterize subcellular RBP localization. With the burgeoning interest in the roles of RBPs in cancer, neurobiology and development, these resources are invaluable to the broad scientific community. Detailed information about these resources is publicly available at the ENCODE portal (https://www.encodeproject.org/).

**Highlights:**

- Antibodies against 365 unique RBPs successfully immunoprecipitate the RBPs
- Short-hairpin RNAs against 276 unique RBPs confirm the specificity of RBP antibodies
- Antibodies characterize subcellular localization of RBPs
- Antibody and hairpin RNA information are provided at https://www.encodeproject.org/

## Introduction

RNA-binding proteins (RBPs) belong to a diverse class of proteins that are involved in co- and post-transcriptional gene regulation (Glisovic et al., 2008). RBPs interact with RNA to form ribonucleoprotein complexes (RNPs), governing the maturation of their target RNA substrates, such as splicing, editing, cap and 3’ end modifications, localization, turnover and translation. Dysregulation of and mutations in RBPs are major causes of genetic diseases such as neurological disorders (Kao et al., 2010; King et al., 2012; Lagier-Tourenne et al., 2010; Nussbacher et al., 2015; Paronetto et al., 2007) as well as cancer (Lukong et al., 2008; Martini et al., 2002; Paronetto et al., 2007). Traditionally, RBPs were identified by affinity purification of single proteins (Sonenberg et al., 1979a; Sonenberg et al., 1979b). Recent advancements in high throughput techniques have identified hundreds of proteins that interact with polyadenylated mRNA in human and mouse cell lines (Baltz et al., 2012; Castello et al., 2012; Kwon et al., 2013).

Genome-wide studies that apply methods such as RNA immunoprecipitation (RIP) (Sephton et al., 2011; Zhao et al., 2010) and crosslinking and immunoprecipitation (CLIP) (Hafner et al., 2010; Konig et al., 2010; Licatalosi et al., 2008; Yeo et al., 2009), followed by high-throughput sequencing (-seq) have identified hundreds to thousands of protein-RNA interaction sites in the transcriptome for dozens of individual RBPs. These sites or clusters have revealed new rules for how RBPs affect RNA processing and novel pathways for understanding development and disease (Hoell et al., 2011; Modic et al., 2013; Wilbert et al., 2012). The availability of antibodies that specifically recognize the RBP and enable efficient immunoprecipitation of the protein-RNA complex is critical for the successful application of these large-scale techniques in a wide range of tissues and cell-types. Alternatively, expression of a fusion protein of one or more peptide tags such as V5, FLAG or HA in frame with the open reading frame of the RBP is also routinely used (Hafner et al., 2010; Wilbert et al., 2012; Zhao et al., 2010), but has several practical and scientific disadvantages. First, it precludes studying the endogenous proteins in human tissues and currently available animal models of disease. Second, creating cell lines that stably express the tagged RBP is labor intensive and has to be performed for every RBP and cell type under investigation. Third, the tags might interfere severely with protein function or target recognition. Lastly, ectopic expression of tagged RBPs typically uses ubiquitously expressed promoters to drive expression, which might alter the endogenous stoichiometry of the RBP to its binding targets. Overexpression in general may complicate the interpretation of results in an irrelevant cell type.

Given these limitations, characterizing antibodies that can specifically enrich for a given RBP is a laborious yet necessary first step for the systematic evaluation of the endogenous RNA substrates of RBPs. In this study we obtained 700 commercially available antibodies that were predicted to recognize 535 candidate RBPs and screened each of them for their ability to efficiently and specifically IP the target RBP. For 51% of the RBPs, we have also identified shRNA reagents that efficiently deplete the target mRNA and protein, simultaneously validating the specificity of the antibodies and providing additional validated experimental reagents. Finally, these antibodies were also used in immunofluorescence assays to determine the subcellular localization of the protein. We expect that this catalog of validated antibodies and shRNA constructs will provide a critical resource for the scientific community.

## Results and Discussion

### A Comprehensive Human RNA Binding Protein Reagent Resource

To comprehensively characterize the protein-RNA interactions and functions of all human RBPs, it is essential to develop a resource of validated antibodies and shRNAs for each RBP. Each antibody must be validated to demonstrate that it efficiently and specifically immunoprecipitates the intended target protein. The efficiency of enriching for the target protein is measured by performing immunoprecipitation followed by western blotting, while the specificity of the antibody is measured by performing shRNA knockdown experiments followed by western blotting. These validated antibodies and shRNAs can also be used in a variety of experiments including CLIP-seq to characterize the transcriptome-wide protein-RNA interactions, immuno-fluorescence studies to characterize the subcellular localization patterns of each RBP, and shRNA knockdowns followed by RNA-seq to characterize the function of each RBP in RNA metabolism.

We have compiled a list of candidate human RBPs from the PFAM database (http://pfam.sanger.ac.uk/), selecting proteins that contain any of the 86 previously known RNA binding domains (Lunde et al., 2007) (Table S1). This list was further filtered to remove proteins containing RNA-binding domains specific for tRNAs, snoRNAs and rRNAs, with the remaining proteins containing domains predicted to bind pre-mRNA or mRNA sequences. Additional proteins such as UPF1 and MAGOH that do not contain canonical RNA binding motifs were added to the list based on their previously characterized associations with RNA regulation. Our primary list of 476 RBPs was then merged with the 845 candidate mRNA binding proteins identified in HeLa cells using interactome capture (Castello et al., 2012). Half of the RBPs in our domain-based list overlapped with the interactome-captured RBPs. The union of these two lists yielded a final list of 1,072 candidate RBPs (Table S1), which for the remainder of this manuscript, will be referred to as the ‘RBP compilation’.

To begin building a human RBP resource, we acquired antibodies interrogating the RBP compilation from Bethyl Laboratories (330 antibodies), MBL International (129 antibodies) and GeneTex (414 antibodies), largely consisting of rabbit polyclonal antibodies. Details about the antibodies including catalog number, host species and antigen information are listed in Table S2. We also acquired 1,139 pre-made shRNA vectors for 491 RBPs from The RNAi Consortium (TRC). The shRNA TRCN ID numbers and the RBP target genes are listed in Table S3. Below we describe our efforts to validate the antibody and shRNA reagents.

### Immunoprecipitation-Western blotting (IP-WB) validation of antibodies

To date, we have tested 700 antibodies, intended to recognize 535 unique RBPs by immunoprecipitation followed by western blotting (IP-WB) validation experiments using K562 whole cell lysate. We utilized an IP protocol that contains a series of stringent washing steps similar to that used in the CLIP protocol. We devised a scheme to score antibodies for their specificity and IP efficiency as outlined in Figure 1, which are largely based on ENCODE ChIP-seq guidelines (Landt et al., 2012). These scores are based on several criteria including the efficiency of IP, the apparent molecular weight (MW) of the target protein (based on the predicted MWs from Genecards (http://www.genecards.org/)), and the number of proteins recognized by the antibody (Figure 1B). The highest quality antibodies are given a score of 1, intermediate quality antibodies are scored 0.5 and low, or unacceptable antibodies, are given a score of 0. In addition, if the protein recognized by the antibody is detected in the immunoprecipitation lane and not detected in the input lane due to expression and/or detection level thresholds, the indicator “IP” is appended to the score (e.g., 1IP). Similarly, if only one protein is detected in the input and immunoprecipitation lanes and is enriched upon immunoprecipitation, but the observed molecular weight deviates more than 20% from the expected molecular weight, the identifier “MW” is appended to the score (e.g., 1MW). Finally, if multiple proteins are detected in the input lane and/or are also enriched upon immunoprecipitation, the identifier “MB” is appended to the score (e.g., 1MB).

**Figure 1.**
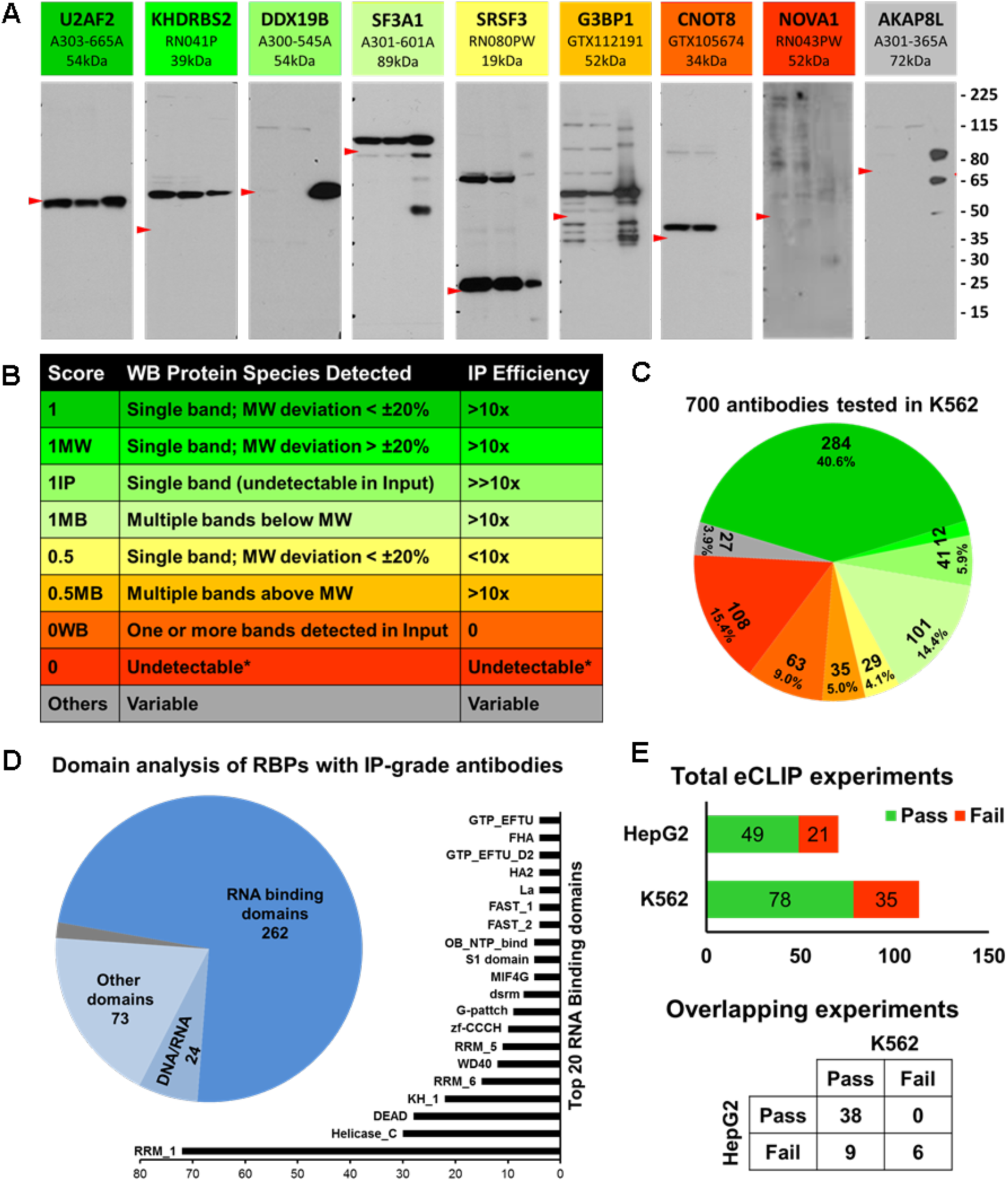
Immunoprecipitation-western blot validation of antibodies against RBPs. (A) Representative blots of antibodies with distinct IP scores. The shades of colors from green to yellow to red represent high, medium and unacceptable quality of antibodies. Grey antibody represents antibodies with ambiguous IP scores. Each blot contains name of the RBP, catalog number and expected MW at the top. Within each blot, lane 1 is 2.5% of input of K562 whole cell lysate, lane 2 is 2.5% of supernatant after IP and lane 3 is 50% of IP pull down sample. Red arrowhead in each panel points to the expected MW of the RBP and size marker (in kDa) is at the right. (B) Table briefly describes the scoring schema with rows of matching colors representing antibodies as in panel A. Column 1 is the IP score, column 2 is the description of protein species detected in the WB and column 3 is IP efficiency deduced from the ratio of band intensities of input and IP pull down lanes. (C) Distribution of IP scores of 700 antibodies validated in K562 cells. Color pattern is consistent with representative antibodies in panel A and score description in panel B. (D) Domain analysis of 365 unique RBPs that have IP-grade antibodies. Pie chart shows the numbers of RBPs with canonical RNA binding domains, putative DNA/RNA binding domains and domains with various other functions. The bar chart shows the top 20 RNA Binding Domains represented in the RBPs. (E) Summary of IP-WB results of eCLIP experiments done to date. Bar chart in the top panel summarizes the total number of antibodies that had either passed or failed the QC step in K562 and HepG2 cells individually. Punnett square at the bottom panel describes the QC results of 53 overlapping eCLIP experiments between K562 and HepG2 cell types.

We identified 284 antibodies (40.6% of tested products) that were characterized with an IP score of ‘1’ (represented by U2AF2 in Figure 1A) indicating that they recognize and enrich only one protein during IP, within 20% of the predicted MW. We identified 12 antibodies (1.7%) that recognize only one protein, but for which the size deviated by more than 20% from the predicted MW, and were therefore scored as ‘1MW’ (KHDRSB2 in Figure 1A). For these antibodies, secondary validations by shRNA depletion of the protein followed by western blotting are necessary to confirm the specificity of each antibody.

We identified 41 antibodies (5.9% of tested products) that were scored as ‘1IP’ in K562 cells indicating that the target proteins were not readily detectable in the input (for example, DHX19B in Figure 1A), but were nevertheless enriched upon IP. The secondary validations to assess the specificity of these antibodies by depletion of the target protein cannot be performed in K562 cells as the protein level in untreated cells is below the detection level of Western blot analysis. However, these antibodies can be further validated for specificity in a different cell line that expresses the protein at a detectable level or by analyzing the whole proteome of the immunoprecipitate by mass spectrometry analysis. For example, antibody A300-864A, which recognizes RBFOX2, was scored as ‘1IP’ in validations using K562 cells, which do not robustly express RBFOX2, but scored ‘1’ when the validation experiment was performed in HepG2 cells which expresses RBFOX2 at a detectable level (Figure S1). 101 (14.4%) of the antibodies recognized multiple proteins *below* the expected MW in the input, supernatant and/or enriched upon IP and were scored as ‘1MB’ (SF3A1 in Figure 1A). On the other hand, 35 (5.0%) of the antibodies recognized multiple proteins *above* the predicted MW and were scored as ‘0.5MB’ (G3BP1 in Figure 1A). Antibodies scored as ‘1MB’ may be suitable for CLIP experiments after passing the secondary validation experiments, as the CLIP-seq protocol involves a size selection step for selecting RNP complexes above the MW of the RBP. CLIP experiments using these antibodies must validate binding events by further downstream analyses. However, due to non-specific bands above the predicted MW, antibodies with a ‘0.5MB’ score cannot be used for CLIP-seq. Because there is no size selection for the RNP complex after immunoprecipitation in the technique, 1MB and 0.5MB products should not be used in RIP-seq experiments, but are nonetheless useful reagents for western blotting.

We scored 29 (4.1%) antibodies as ‘0.5’ (Low Enrichment-Low Priority), for which we observed that the efficiency of immunoprecipitation is relatively low such that the intensity of the bands detected in 50% of the immunoprecipitate (Lane 3, SRSF3 in Figure 1A) is less than the band intensity of 2.5% Input sample (Lane 1). An additional 3.9% of the tested antibodies represented by AKAP8L in Figure 1A, had multiple complicating criteria including multiple bands (MB), MW discrepancy (MW) and detected only upon enrichment (IP) and were designated as ‘1MBMW’, ‘1IPMW’ etc. Due to ambiguity in the specificity of these antibodies, these are considered low priority (‘Others’ in the Table S4). Finally, there are 171 (24.4%) antibodies that were scored as failing IP validation K562 cells, because they either do not recognize the correct protein or do not enrich the target protein upon immunoprecipitation. These antibodies were further grouped into two categories. Antibodies that recognize the correct protein in the input lane, but failed to enrich the protein in IP are scored as ‘0WB’, like GTX105674 recognizing CNOT8 in Figure 1A that can only be used for western blotting (63, 9.0%). Additionally there are 108 (15.4%) antibodies that neither recognized nor enriched the correct protein in K562 cells (RN043PW against NOVA1 in Figure 1A). This failure might be due to either the protein recognized by the antibody is not expressed in K562 cells or the antibody is simply ineffective for IP and/or WB. The cell type specificity could in the future be evaluated by validating the antibody in other cell types known to express the RBP.

The results of the IP-WB validations performed to date are summarized in Figure 1C and in Table S4, which includes 438 (62.5%) antibodies against 365 unique RBPs that scored 1, 1MW, 1IP and 1MB and thus are categorized as ‘IP-grade’ based on our protocol and scoring criteria in the indicated cell type. Diversity of these 365 RBPs was analyzed by searching for Pfam domains (pfam.xfam.org) associate with these proteins. Biomart tool (http://www.ensembl.org/biomart/) was used to search for ENSEMBL and Pfam IDs using Uniprot IDs as query. If a RBP has multiple copies of same Pfam domain then that ID was counted only once for that RBP and if a RBP has more than one type of Pfam domain all of them were counted. This analysis identified 322 Pfam domains associated with 359 RBPs with 680 total occurrences (see Table S5). 322 Pfam domains are classified into three groups. 159 domains are either directly bind RNA or are associated with RNA processing and 268 RBPs having at least one of these domains are considered as true RBPs (Figure 1D, pie chart). There are 34 Pfam domains that are either putative DNA/RNA binding domains or bind DNA/chromatin to regulate transcription that occur in 23 RBPs. Another 68 RBPs contain 139 domains that have no direct role in RNA/DNA binding or RNA processing or no known function. Among the 268 true RBPs, RRM_1 domain (PF00076) is the most frequently found domain present in 72 RBPs followed by Helicase_C and DEAD domains. The top 20 domains present in the 268 RBPs are shown in Figure 1D bar chart.

All of IP-grade antibodies have been or are in the process of being further validation by shRNA knockdown experiments as described in the next section. IP-WB images for all 700 antibodies characterized in K562 cells are available at the ENCODE project portal (https://www.encodeproject.org/) and can be identified using the ENCAB accession IDs as explained in Box 1 and Figure 2. As additional antibody validation experiments are performed, the results will also be added to the ENCODE project portal under the same accession IDs.

**Figure 2.**
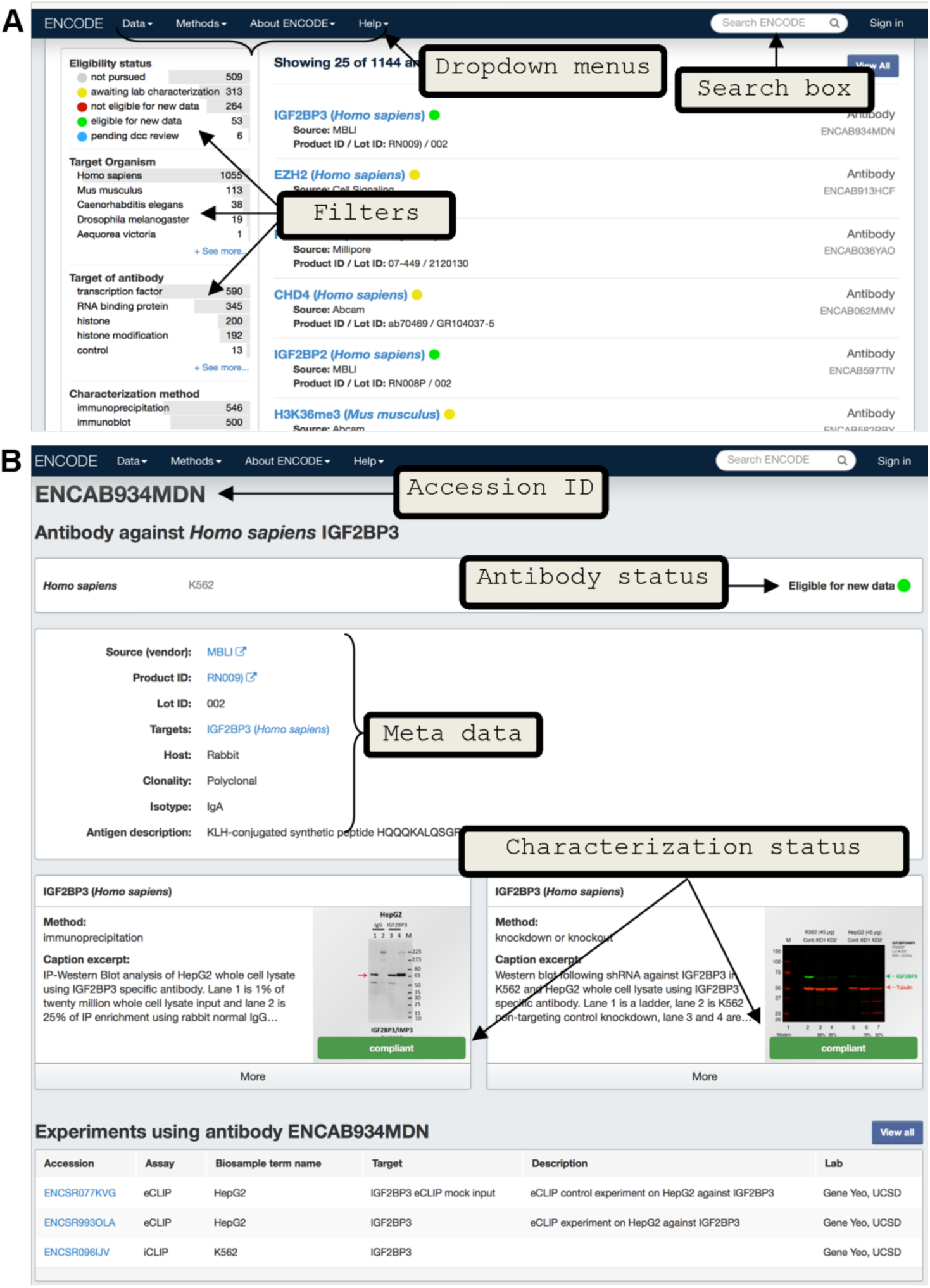
Accessing antibody characterizations in ENCODE portal. (A) Screen shot of the ‘Antibodies’ page of the ENCODE portal. Arrows point to dropdown menus at the top and filtering criteria (to narrow the search) in the left side of the page. (B) Screen shot of a representative antibody characterization page from the portal. Top of the page contains the ENCAB accession ID and antibody status. Antibody metadata like catalog number, link to vendor page, lot number and other information are listed in the top middle panel. Characterizations are in the middle of the page which can be expanded by clicking ‘more’ option. These characterization subpages lists the submitter lab name, link to download image file and ENCODE standards document which was used to review the characterization. Links to any experiments that have used the antibody are also listed at the bottom of the page.

As part of the ENCODE project, we are performing CLIP-seq experiments using our eCLIP method (Van Nostrand EL et al, manuscript under preparation). The IP-WB images that are generated during eCLIP experiments have an additional lane of IP using host-species matched normal IgG antibody as control and can be accessed using the same ENCAB IDs in the portal. We observe that only 70% of our *IP-grade* antibodies pass this internal QC step during eCLIP experiments, presumably due to differences in the IP protocol initially used and that used in the eCLIP protocol. We used a protocol similar to the IP steps in the iCLIP method (Konig et al., 2010; Konig et al., 2011) for our initial 700 experiments in K562 cells that included an overnight pull down step (see supplementary methods for details), whereas in our eCLIP protocol, the RNP complexes are immunoprecipitated for only 2 hours at 4°C (Van Nostrand EL et al, manuscript under preparation). Besides protocol differences, the possibility of antibodies going bad during two year storage period also can’t be excluded. Nonetheless, of the 113 eCLIP experiments attempted in K562 to date, 69% passed the IP-WB QC step (Figure 1D). We presume to generate about 300 eCLIP experiments using our 438 IP-grade antibodies.

As cell-type specific expression and post-translational modifications add additional layers of complexity affecting the success of immunoprecipitation of RBP-RNA complexes, we tested the utility of IP-grade antibodies validated in K562 cells for eCLIP experiments in HepG2 cells. Of the 53 RBPs that were subjected to eCLIP experiments in both K562 and HepG2 cells, 72% (38) of them passed the QC step in both cells types; 9 antibodies failed the IP-WB step in HepG2 cells but passed in K562 cells and another 6 of them failed in both cell types (Figure 1E). Of the 70 eCLIP experiments, which includes the 53, performed in HepG2 cells, 70% (49) passed the QC step, which is comparable to the success rate in K562 cells. Thus we conclude that the majority of antibodies that are considered *IP-grade* by our criteria in K562 are likely to work in HepG2 cells. If an eCLIP experiment was attempted in HepG2 cell line, the portal will also have IP-WB images from the HepG2 experiments under same accession ID.

### Secondary antibody validation by short hairpin RNA transduction

To verify the target specificities of these antibodies, we performed secondary validations using shRNA-mediated RNA interference. Specifically, to conclude that an antibody recognizes the intended protein, and not a different protein that migrates at the same size range as the intended protein, the band identified in the IP-Western blot must be decreased by at least 50% in shRNA knockdown cells compared to cells expressing a control shRNA. To do this, we first identified 1,139 shRNAs from the RNAi Consortium (TRC) that target 491 RBPs (Table S3). To date, we have tested a total of 370 shRNAs against 273 unique RBPs. 274 shRNAs against 242 unique RBPs have been tested in K562 cells and 333 shRNAs against 265 unique RBPs have been tested in HepG2 cells. Of these, 237 shRNAs against 234 RBPs have been tested in both cell lines. We defined a successful knockdown as shRNAs resulting in >50% reduction of the target mRNA or protein, compared to control cells transduced with a non-target control (NTC) shRNA, depending on whether depletion is monitored by qRT-PCR or western blotting. Of the 274 shRNAs tested in K562 cells, 70% passed the validation criteria by RT-qPCR and 60% passed the western blotting validation (Figure 3A). Similarly, in HepG2 cells 62% and 55% of the target mRNAs and proteins were depleted >50%, respectively, as monitored by qRT-PCR or western blotting (Figure 3B). In most cases, we observed reasonable correlation between the extents of depletion of both the mRNA and protein between both cell lines, though overall, the protein depletion efficiency is an average of 1.25-fold greater in HepG2 cells than in K562 cells (median depletion efficiency of 72% vs. 68%) (Figure 3C, D). Overall, 68.4% of RBPs tested in both cells were depleted at the protein level more than 50% in both K562 and HepG2 cells while 21% of RBPs are depleted >50% in one cell but <50% in the other cell type (Figure 3D).

**Figure 3.**
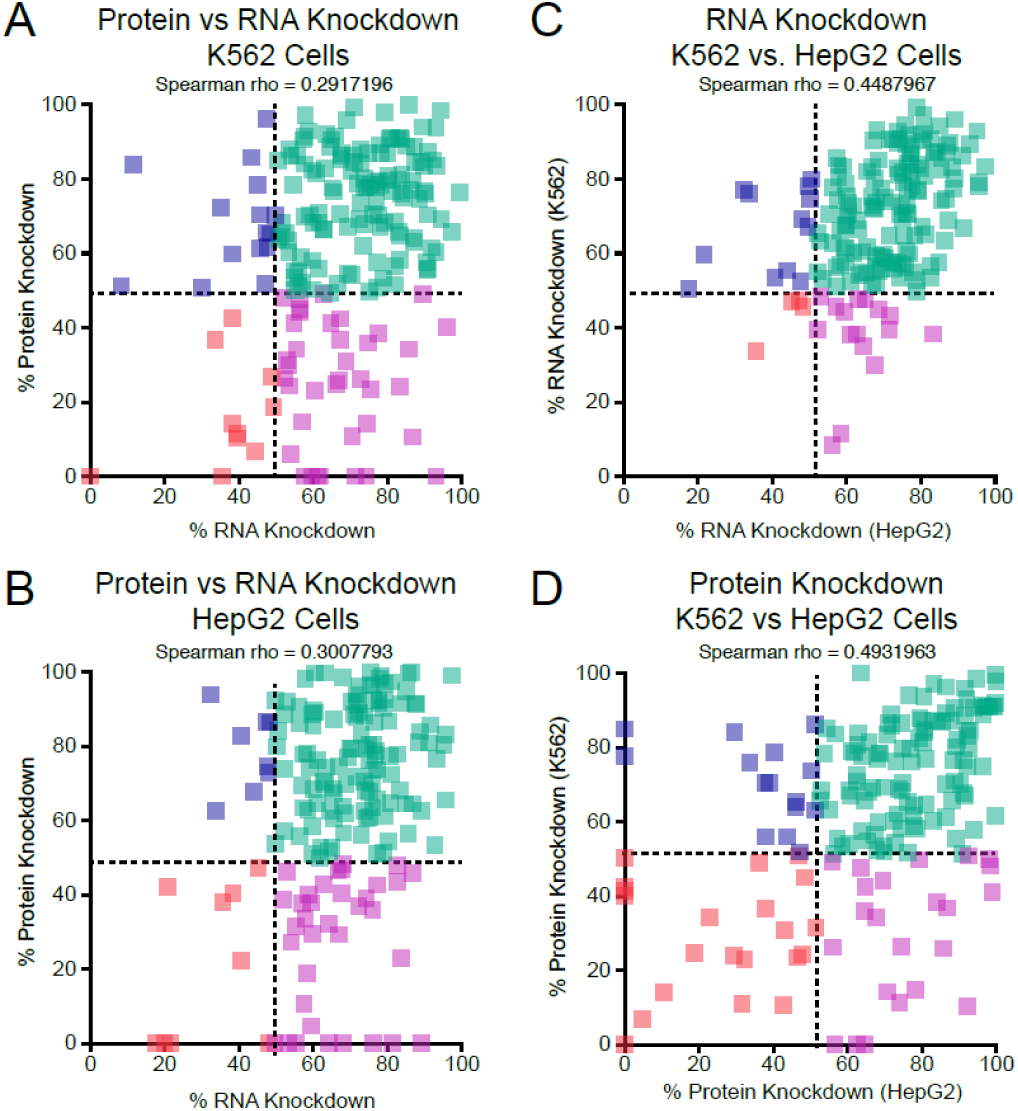
Comparison of mRNA and protein depletion in shRNA knockdown experiments. (A) Comparison of protein and RNA knockdown efficiency in K562 cells. (B) Comparison of protein and RNA knockdown efficiency in HepG2 cells. (C) Comparison of RNA knockdown efficiency between K562 and HepG2 cells. (D) Comparison of protein knockdown efficiency between K562 and HepG2 cells.

Of the 284 antibodies that were scored as ‘1’ during IP validation, 183 were tested by KD-WB in K562 cells and 184 were tested in HepG2 cells. Many of these antibodies, as exemplified by the antibody recognizing PABPC4 (Figure 4A), recognize only a single protein in the control shRNA-treated sample lane and the band intensity was reduced >50% in the RBP shRNA knockdown lanes. 74 of the antibodies that were scored as ‘1’ during IP validation were found to recognize multiple bands in the KD-WB experiments in both the control and RBP shRNA lanes, but only the band of the predicted MW was depleted >50% in the RBP shRNA knockdown lanes. The antibody that recognizes KHSRP is example of such a case and recognizes proteins of ~40 kDa and ~80 kDa, but only the ~80 kDa band, near the predicted MW of KHSRP of 73 kDa, is depleted in the KHSRP shRNA samples (Figure 4B). We presume that the difference in the reagents (e.g. secondary antibodies) and detection sensitivity between chemiluminescence used for the IP-WB experiments and fluorescence used for the KD-WB experiments are most likely the source of different band patterns observed between the experiments. In these cases, the multiple bands detected by fluorescence are likely to be background or cross-reactivity, because only one band was detected when the same sample was analyzed using chemiluminescence. For the four antibodies we have tested so far which detected and immunoprecipitated a protein with an aberrant MW in the IP validation experiments (and received a score of 1MW), we also detected a band with the same aberrant MW in the control sample of KD-WB. In each case, the band with the aberrant MW was depleted upon shRNA knockdown, confirming the identity of the protein detected by the antibody. For example, the predicted MW of FASTKD2 is 81 kDa, yet the FASTKD2 antibody recognizes a protein of ~60kDa in both K562 and HepG2 cells (Figure 4C). The protein band recognized by FASTKD2 antibody is likely to either be alternative protein isoform or post-translationally modified form of FASTKD2. For 6 of the antibodies we have tested by KD-WB that detected multiple bands (1MB) in the IP-WB validation, we also detected multiple bands in the control shRNA sample of KD-WB. For five of these, the band closest to the predicted MW was depleted >50% in the RBP shRNA sample, and the intensity of most other bands are comparable between the control and RBP shRNA lanes, indicating that the target RBP is one among the multiple bands detected by the antibody. Some cases are additionally complicated. For example, the ADAR antibody scored a 1 MB in the IP-WB experiments, recognizes multiple bands in the KD-WB experiments, and more than one band is observed to be depleted in the ADAR1 shRNA samples (Figure 4D). However, ADAR1 is annotated as expressing multiple protein isoforms and it is likely that the antibody recognizes more than one ADAR1 isoform. The specificity of ‘1MB’ antibodies, which have multiple bands in the IP enrichment that are not depleted via shRNA knockdown, are not considered to be fully validated and therefore should not be used for CLIP experiments. The KD-WB data images can be publicly accessed through the ENCODE DCC portal (https://www.encodeproject.org/) and up-to-date report on the status of the RBP antibody validation experiments can be obtained at https://goo.gl/pZqDR5. Table S6 summarizes the results of the RT-PCR and KD-WB experiment for 370 shRNA constructs and also contains the TRC number of the shRNA plasmids, the target sequence of the shRNA, sequences of primers used for RT-qPCR validation, and catalog numbers of antibodies used in the KD-WB characterization.

**Figure 4.**
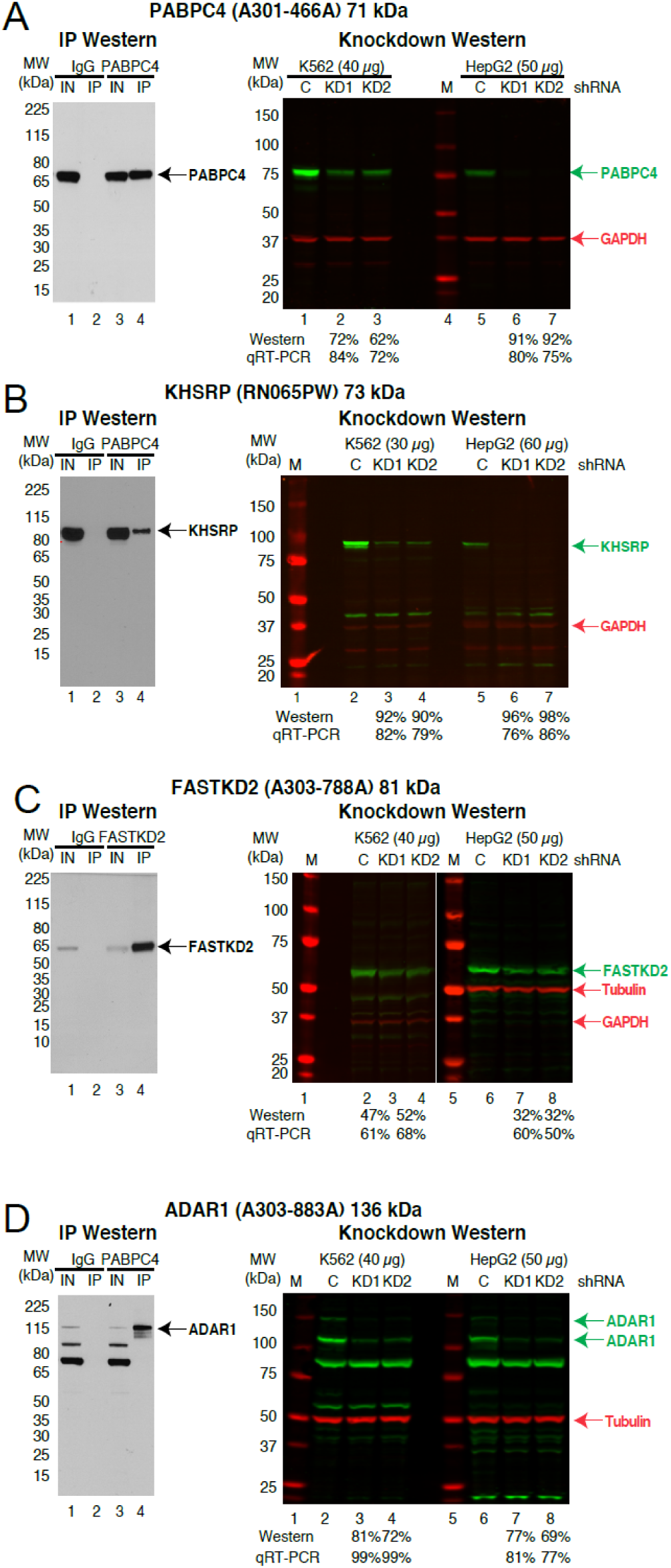
shRNA knockdown-western blot validation of antibodies against RBPs. Representative IP-WB (left) and shRNA knockdown-western blot validations (right) for antibodies that cover a spectrum of band patterns. Experiments are shown for antibodies that recognize PABPC4 (A), KHSRP (B), FASTKD2 (C), and ADAR1 (D). For each experiment the molecular weight markers are shown along with the percent depletion in the knockdown sample compared to the control shRNA sample for both the western blot and qRT-PCR experiments. The position of the RBP (green) and the loading controls of GAPDH or Tubulin (red) are shown.

### Immuno-Labeling Studies

As an additional level of validation, and to gain broader biological insights into RBP function, we have conducted immuno-fluorescence (IF) studies using the RBP compilation antibodies in conjunction with different subcellular markers. We observed clear subcellular distribution patterns for the majority of the antibodies tested in HepG2 cells. These results are generally consistent with the known subcellular localization features of previously characterized RBPs. For example, the DDX21, BUD13 and GRSF1 proteins are respectively localized to nucleoli, nuclear speckles and mitochondria (Figure 5), consistent with the known functions of these RBPs in rRNA maturation (Calo et al., 2015), splicing control (Zhou et al., 2013) and mitochondrial biogenesis (Antonicka et al., 2013; Jourdain et al., 2013). The full repertoire of results obtained through these systematic imaging studies have been organized within a resource imaging database that will be described in more detail in a separate manuscript (Lécuyer et al, in preparation).

**Figure 5.**
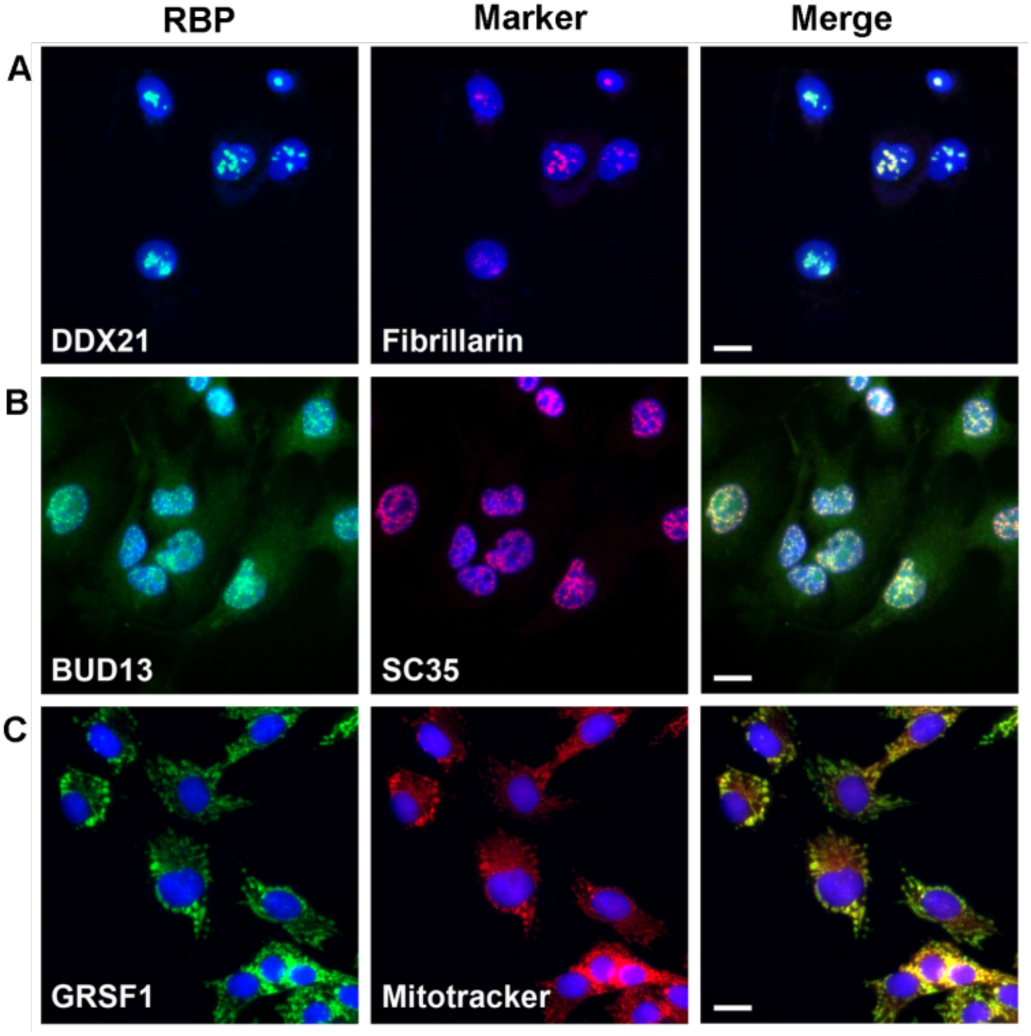
Immunofluorescence characterization of antibodies. (A-C) Representative images of immunofluorescence characterizations of antibodies. Left column is the images of RBPs pseudo-colored in green, center column is the sub-cellular markers pseudo-colored in red and the right column is the merged image of RBP, subcellular marker and nuclear stain (blue). Scale bar in the merged image represents 20nm. (A) DDX21 antibody (RN090PW) co-stained with nucleolar marker fibrillarin. (B) BUD13 antibody (A303-320A) co-stained with nuclear speckles marker SC35 and (C) GRSF1 antibody (RN050PW) costained with mitochondrial marker mitotracker.

In conclusion, we have comprehensively validated antibodies and shRNA constructs for hundreds of unique human RBPs. The scoring schema described for the IP-WB validations can be extended to future large-scale antibody characterization studies. The publicly accessible reagent collections serve as key resources for the illumination of functional RNA elements in the human transcriptome.

## Experimental procedures

### Immunoprecipitation-western blot validation

Five million human K562 cells were lysed, sonicated (instead of DNase treatment) and the whole cell lysate used for IP characterizations. Five micrograms of antibody coated on Dynabeads (coupled with either Protein A or anti-Rabbit IgG or anti-mouse IgG) was used to IP overnight at 4°C. Protein-enriched beads were washed twice with a high salt wash buffer containing 1M NaCl and detergents to reduce non-specific interactions. For the western blot analysis, aliquots of the input (2.5%), supernatant (2.5%) and bound fractions (50%) ran on 4-12% SDS-PAGE gel and transferred onto PVDF membrane. Membrane was incubated with 0.2-0.5 μg/ml of the same antibody used for IP as the primary antibody. TrueBlot HRP secondary antibodies were used to avoid IgG heavy and light chain immunoreactivity. See the supplementary methods for detailed protocol.

### shRNA knockdown-western blot validation

The shRNA constructs are in pLKO plasmids to facilitate the production of lentiviral particles following co-transfection with appropriate packaging vectors in 293T cells. Lentiviral particles were tittered by qPCR and used to transduce 0.5-0.7 million K562 cells or 0.5 million HepG2 cells in biological duplicate at a MOI of 10. One day after transduction, puromycin was added to the media and the cells were subjected to selection for 5-6 days after which we harvested both RNA and protein (see supplementary methods for detailed protocol).

### Image database

Western blot images of both IP and shRNA knockdown experiments are uploaded on to ENCODE portal which is built and maintained by the Data Coordination Center (DCC). See the Box 1 for description about how to access the database.

### Box 1: Accessing the ENCODE portal

Antibody characterizations can be accessed in the ENCODE portal (www.encodeproject.org/) (Solan et al., 2015) (Figure 2A). The text search box on the top right of the page can be used to search for a particular RBP of interest as well as an antibody or shRNA construct using their catalog or TRC numbers respectively. Alternatively, the user can browse the entire collection of antibodies by opening the drop-down menu ‘Data’ on the top of the page and then choosing ‘Antibodies’. In the Data/Antibodies page, the results can be filtered using options such as ‘Eligibility status’, ‘Target of antibody’, ‘Characterization method’, ‘Source’, ‘Lab’ etc. Other than the ‘Eligibility status’, other filtering criteria are straight forward for first time users (https://www.encodeproject.org/help/antibody_characterization_process/). The Eligibility status categorizes antibodies as ‘not pursued’, ‘awaiting lab characterization’, ‘eligible for new data’ and ‘not eligible for new data’ based on whether the characterizations met ENCODE standards. Each antibody is rigorously reviewed by the Data Coordinating Center (DCC), a subgroup in ENCODE consortium independent of the labs that characterized the antibodies, according to the ENCODE Standards document (https://www.encodeproject.org/about/experiment-guidelines/#antibody; (Landt et al., 2012) and assigned to one of the four eligibility statuses described below.

**‘not pursued’:** Indicates that the lab has planned to characterize the antibody and may have done preliminary primary characterizations in one cell type but it is not interested in doing further characterizations at present. Antibodies under this category are open for further validation either in the same or different cell type.

**‘awaiting lab characterization’:** Indicates that the lab has completed a primary characterization in one cell type and is expecting to do further validations.

**‘eligible for new data’:** Indicates that both primary and secondary characterizations are performed in at least one cell type, both of which met the ENCODE standards. Antibodies under this category are eligible for new data generation in the same cell type. A primary validation in the second cell type is necessary before generating new data in that cell type.

**‘not eligible for new data’:** Indicates that both primary and secondary validations are done in one or more cell types but they did not meet the ENCODE standards. These antibodies are not eligible for new data generation.

Each unique pair of antibody catalog and lot numbers is given an ENCODE accession number (ENCABnnnxxx). An antibody can be selected by clicking the RBP name listed in the Data/Antibodies page, directing the user to an ‘antibodies/ENCABnnnxxx’ page (Figure 2B). The ENCABnnnxxx page contains information regarding species and cell type in which that antibody was characterized, antibody metadata (vendor, host, antigen etc) and the characterizations. The characterization sub-page is expandable by clicking the ‘more’ option, which lists further information such as characterization methods (immunoprecipitation, knockdown-WB etc), image caption, submitter and lab names as well as links to download the characterization image and the version controlled standards documents used to set the eligibility status. Each antibody characterization is also classified into ‘compliant’, ‘not compliant’ or ‘not submitted for review by lab’ which indicate that they met, did not meet the standards or were not reviewed based on the standards document. These characterization statuses determine the eligibility status of the antibody. For example all antibodies with ‘eligible for new data’ status are expected to have both primary and secondary characterizations with ‘compliant’ status. For the antibodies with ‘eligible for new data’ status, the page also will contain links to experiments (eCLIP, for example) in which that antibody might have been used. For more information about data available at the ENCODE Portal, please see the Getting Started help page (https://www.encodeproject.org/help/getting-started/) and Solan et al., 2015.

## Acknowledgement

We thank B Williams and J Fahrni of Bethyl Labs, S Kendall and B Parmakhtiar of GeneTex and S Kitamura of MBLI for their support in antibody collections. We also extend our thanks to the members of Yeo and Graveley laboratories for their critical comments on the manuscript. This study is funded by NIH grant HG007005 to BRG and GWY and NIH grants HG004659, NS075449 to GWY. GWY is an Alfred P. Sloan Research Fellow.

---

**Figure S1.**
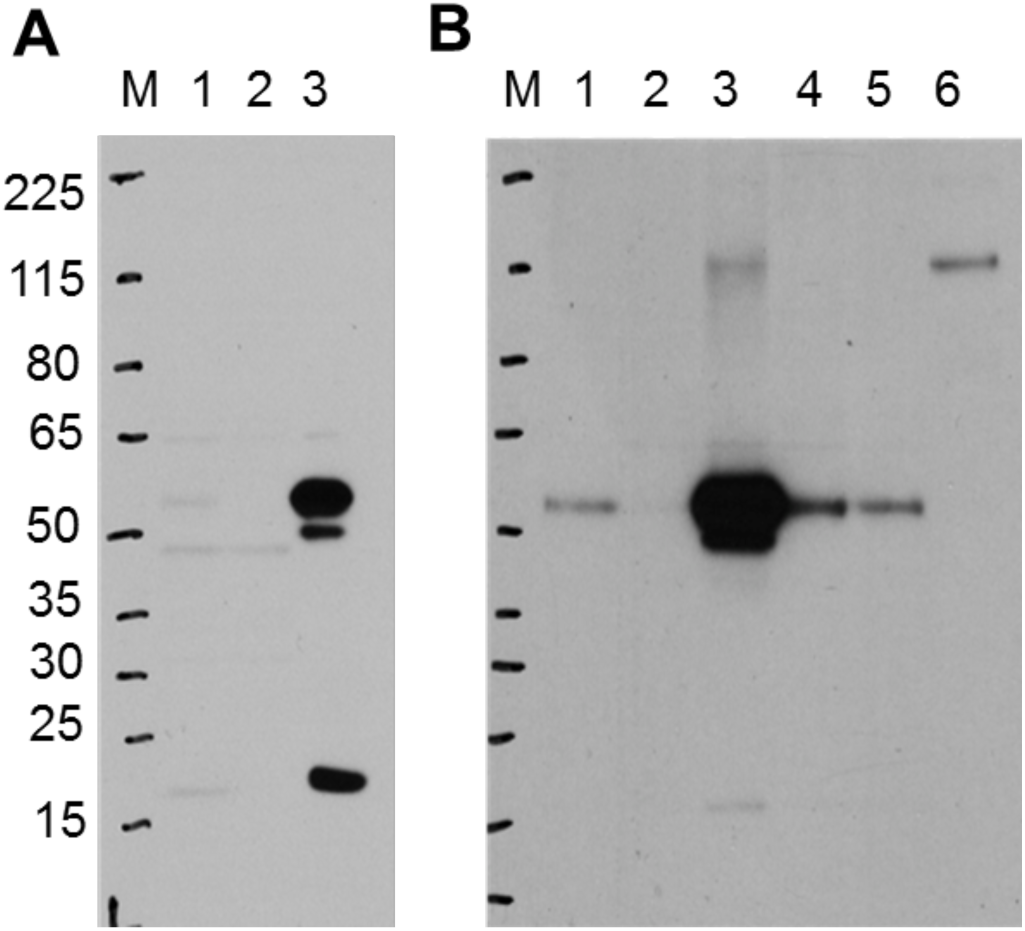
Antibody IP-scores are cell type dependent. Both panels A and B are IP validation of A303–864A antibody against RBFOX2 protein. (A) IP validation using K562 whole cell lysate. The validation was scored as ‘1IP’ as the protein band could not be detected in the Input lane. (B) IP validation using HepG2 whole cell lysate and the validation was scored as ‘1’. In both (A) and (B), lane 1 is 2.5% whole cell lysate, lane 2 is 2.5% of supernatant after IP and lane 3 is 50% of IP pull down using RBFOX2 antibody. Lanes 4–6 are the same but using Rabbit normal IgG.

## Supplementary Tables

**Table S1. RBP Compilation**

Sheet 1 is the list of 476 proteins that are compiled based on presence of the RNA Binding Domains (RBDs).

Sheet 2 is the ‘1072 RBP Compilation’ list which is the union of 476 RBPs identified by RBDs and the 845 proteins identified in the RBP interactome capture study (Castello et al., 2012). Sheet 3 is the list of ENSEMBL IDs for the 1072 RBP Compilation and associate Pfam domain IDs resulted from Biomart tool (http://www.ensembl.org/biomart/) search.

**Table S2. RBP Antibody Details**

Sheet 1, 2 and 3 contain metadata like host species, clonality, antigen information, purification method etc for antibodies from Bethyl, MBLI and GeneTex companies respectively. Each sheet may vary in number of columns and specific information based on disclosure policy of the vendors.

**Table S3. RBP shRNA Details.**

Information about all RBP shRNAs acquired including the target genes.

**Table S4. Results of the IP-WB validations.**

Sheet 1 is the list of all 700 IP-WB experiments performed using K562 cells, containing catalog and lot numbers of antibodies, Uniprot ID and expected MW of RBPs, IP score, ENCAB ID and antibody status for each antibody.

Sheet 2 is distribution for 700 antibodies by IP score as well as by vendor wise breakdown. Sheet 3 is the Uniprot IDs for 535 unique RBPs for which at least one antibody was tested in K562.

Sheet 4 is the list of 438 IP-grade antibodies.

**Table S5. Domain analysis of 365 unique RBPs having IP-grade antibodies.**

Sheet 1 is the list of ENSEMBL, HGNC symbol and Pfam IDs for 365 unique RBPs that have IP-grade antibodies.

Sheet 2 is the list of 332 Pfam IDs associated with 365 RBPs.

Sheet 3 is the total occurrence of each Pfam domain types among 365 RBPs.

**Table S6. Summary of the shRNA-KD validation experiments for 370 shRNA constructs.**

Information presented includes the TRC number of shRNA plasmids, sequences of primers used for RT-qPCR validation and catalog numbers of antibodies used in the KD-WB characterization.

## Supplemental Methods

### Immunoprecipitation followed by Western blot (IP-WB)

K562 cells (ATCC CCL-243, lot 59300853) grown in RPMI 1640 (Life Technologies, 11875119) with 10% FBS (Life Technologies, 26140079) and 1% Pen-Sterp (Life Technologies, 15140163) were flash frozen into 5×10^5^ aliquotes and stored at -80°C. Cell pellets were thawed on ice and resuspended in 450ul Lysis buffer containing 50 mM Tris-HCl, pH 7.4; 100 mM NaCl; 1% NP-40 (IGEPAL CA-50); 0.1% SDS; 0.5% sodium deoxycholate with protease inhibitor (Roche, 11836170001). Cell lysate left in ice for at least 15 minutes for complete cell lysis and then sonicated using Bioruptor for 5min with 30sec on/off pulses in cold water bath. Cell lysate spun at 18000g for 20min in a refrigerated centrifuge and the clear supernatant was stored in new tube on ice for IP. 0.625mg per tube of Dynabeads (anti-mouse (11202D) or anti-rabbit IgG (11204D) secondary or Protein A (10002D) coated beads) washed twice with 900ul cold Lysis buffer and resuspend beads in 250ul of Lysis buffer. 50ul of the washed beads were used to preclear the whole cell lysate by rotating for 30min at RT. With remaining 200ul of beads, 5ug of antibody specific to RBP of interest was added and rotated for at least 1hr at RT for antibody-bead coupling. After preclearing, tubes were kept on magnetic stand (Dynamag, 12321D) and clear lysate transferred to new tube. 25ul (5%) of this precleared lysate was stored at -20°C as *Input sample* for WB later. After one hour of antibody-bead coupling, excess antibodies were removed and beads were washed three times with 900ul cold lysis buffer on magnetic stand. Precleared cell lysate was then added to the antibody coupled beads and rotated at 4°C. After overnight immunoprecipitation at 4°C, tubes were kept on magnetic rack, 25ul (5%) of clear supernatant/ flow through was saved as *IP-sup sample* for WB later. Excess flow through was discarded and beads were washed two times in 900ul cold **High Salt Wash** buffer containing 50 mM Tris-HCl pH 7.4, 1 M NaCl, 1 mM EDTA, 1% NP-40, 0.1% SDS, 0.5% sodium deoxycholate. Beads were then washed two more times with 900ul cold **PNK Wash buffer** containing 20 mM Tris-HCl pH 7.4, 10 mM MgCl2, 0.2% Tween-20. Washed beads were resuspended in 26ul PNK Wash buffer, 10ul 4X LDS sample buffer (Novex, B0008) and 4ul 1M DTT. With the 25ul Input and IP-Sup samples saved previously, 10ul 4X LDS sample buffer and 4ul 1M DTT were added and mixed. All the samples were denatured by boiling at 70°C for 10min with shaking at 1200 rpm (in Thermomixer). After boiling, samples were immediately placed on ice cold magnetic stand and liquid was removed for gel loading. Samples were loaded on appropriate denaturing 15 well, 1.5mm thick, 4–12% Bis-Tris gel (Novex, NP0336BOX) for RBPs with MW less than 225kDa and 8% Tris-Glycine (Novex, EC6018BOX) for RBPs with MW more than 225kDa. Spectra Broad (Thermo, 26634) or High range (Thermo, 26625) MW markers were used depending on MW of RBP. Samples were run at 150V for 1.30hr using MOPS-SDS running buffer (NP0001). Protein bands were then transferred on to PVDF membrane (BioRad, 1620177) using NuPAGE transfer buffer (NP00061) with 10% methanol at 200mA constant current for 2hr in cold room. Transfer efficiency was checked by Ponceau staining and blots were blocked with 5% skim milk (Genesee Scientific, 20–241). Membranes were then incubated overnight at 4°C with 0.2–0.5μg/ml (usually 1:2000–1:5000) of the same antibody used for IP as primary antibody. After primary antibody incubation, membranes were washed three times in 1X TBST for 10min and incubated for 1–3hr at RT with appropriate secondary TrueBlot HRP antibody (Rockland, 18–8816–33, 18–8815–33). After secondary incubation, membranes were washed three times in TBST and developed with ECL (Pierce, 32106) or ECL Plus (Pierce, 32132) substrates depending on the abundance of RBP. Protein marker bands and edges of the membrane were marked down in the photo-developer film which was scanned to generated images. IP results were scored based on the following scoring schema.

**1**: Only one band, that deviates less than 20% from the expected molecular weight and detected in input lane. The same band is also enriched upon immunoprecipitation with band intensity greater than or equal to that in the Input lane.

**1IP**: One band is detected only upon immunoprecipitation enrichment that deviates less than 20% from the expected molecular weight. The same band is NOT detected in the input lane due to expression and/or detection level thresholds.

**1MW**: Only one band is detected in the input and immunoprecipitation lanes, but the observed molecular weight deviates more than 20% from the expected molecular weight. The same band is also enriched upon immunoprecipitation with a band intensity greater than or equal to that in the input lane.

**1MB**: One prominent band is detected in input and immunoprecipitation slanes that deviates less than 20% from the expected molecular weight. This band is also enriched upon immunoprecipitation with band intensity greater than or equal to that in the input lane. In addition, there are multiple bands *below* the expected molecular weight, which are also detected in the input lane, and/or enriched upon immunoprecipitation.

**0.5MB**: One prominent band is detected in input and immunoprecipitation lanes that deviates less than 20% from the expected molecular weight. The band is enriched upon immunoprecipitation with an intensity greater than or equal to that in the Input lane. In addition, there are multiple bands *above* the expected molecular weight that are also detected in the input lane and/or enriched upon immunoprecipitation.

**0.5**: Only one band is detected in input lane and deviates less than 20% from the expected MW. The same band is *poorly* enriched up on immunoprecipitation with band intensity *less than* the input lane.

**0WB**: The antibody does not enrich the protein upon immunoprecipitation but detects band at the expected MW.

**0**: The antibody neither detects bands in input lane nor enriches upon IP using K562 lysate.

### Production of shRNA lentiviral particles

0.8–1×106 293 T cells (catalog number: CRL-11268, ATCC) were plated in each well of 6-well plate with 10 % FBS (catalog number: 30–2020, ATCC), DMEM (catalog number: 11995–065, Life technologies) medium without penicillin and streptomycin. The cells were incubated overnight at 37°C and were typically 70–80% confluent. A cocktail for each transfection was assembled in a polypropylene tube containing 500 ng pLKO-shRNA, 500 ng psPAX2 Packaging DNA, 50 ng PMD2.G Envelope DNA, and serum-free OPTI-MEM to 100 μl. 3.1 μl of FuGENE HD Transfection reagent (Catalog number: E2311, Promega) was added to the tube (FuGENE:DNA=3:1), incubated for 20 minutes at room temperature, the DNA mix was gently added dropwise to the cells, followed by incubation at 37°C for 12–15 hr. The next day, the media was changed to remove the transfection reagent, and the cells were washed with PBS once and 1.5 ml fresh media +10% FBS + penicillin/streptomycin was added. After incubation at 37°C for four days, the media was harvested from cells, stored at 4°C, and 1.5 ml of fresh media was added and incubated overnight at 37°C. The media was harvested and pooled with the media collected on day 4, spun at 1250 rpm for 5 min to remove cells and the viral stock stored at -40°C.

### qRT-PCR Lentivirus Titration Assay

Lentiviral titrations were performed using the qPCR Lentivirus Titration kit from Applied Biological Materials Inc. (Catalog Number LV900). 2 μl of the viral supernatant was added to 18 μl of Virus Lysis buffer and incubated at room temperature for 3 minutes. qRT-PCR reactions were assembled to include 12.5 μl of 2X qRT-PCR Mastermix, 2.5 μl of the viral lysate, and 10 μl of the reagent mix. qRT-PCR was performed by incubation at 42°C for 20 minutes, 95°C for 10 minutes, followed by 40 cycles of 95°C for 15 seconds and 60°C for 1 minute. Viral titers were calculated from the Ct values by using the Applied Biological Materials Inc. on-line lentiviral titer calculator at http://www.abmgood.com/High-Titer-Lentivirus-Calculation.html

### Lentiviral Transduction of K562 and HepG2 cells

K562 cells (ATCC CCL-243 (lot 59300853)) were grown in 500 ml RPMI-1640 with glutamine medium (Hyclone, SH30027.01), 50 ml Fetal Bovine Serum (FBS) (10% Final Concentration) (Hyclone, SH30071.03), and 5 ml Pen-Strep (1% Final Concentration) (Invitrogen, 15140–163). HepG2 cells (ATCC HB8065 (lot 59635738)) were grown in 500 ml DMEM (HyClone, SH30022.01), 50 ml Fetal Bovine Serum (FBS) (10% Final Concentration) (Hyclone, SH30071.03), and 5 ml Pen-Strep (1% Final Concentration) (Life Technologies, 15140122). Frozen stocks of cells were thawed by gentle agitation in a 37°C water bath and the cells into the growth medium and centrifuged at 1000 rpm for 5 minutes. The cell pellets were resuspended in fresh growth medium and incubated at 37°C in a 5% CO2 air atmosphere incubator. The growth medium was changed every 2 to 3 days and the cells split when the cell density reached 70–80% confluence. For transductions, 5×10^5^ cells were plated into each well of 6-well plates and incubated for overnight such that the cells were 50–60% confluent. The media was exchanged with fresh media containing 8 μg/ml of polybrene (Catalog Number H9268, Sigma-Aldrich) and lentiviral particles added to an MOI of ~10. After 24 hours, fresh media containing 3 μg/ml of puromycin was added and the cells incubated for 48 hours followed by a second exchange of media containing puromycin, incubation for 48 hours, a third exchange of media and incubation for 24 hours. On day 6, the cells were harvested and half of the cells used to prepare RNA for qRT-PCR and the other half used to prepare protein lysate for western blotting.

### RNA Isolation

RNA isolation was performed using a Promega Maxwell 16 Instrument and the Maxwell 16 LEV simplyRNA Cells Kits (Catlog Number AS1270). Briefly, cells were pelleted by centrifugation at 300 × g for 3 minutes and the medium removed. 200 μl of chilled 1-Thioglycerol/Homogenization solution was added to the cell pellet and vortex until dispersed. 200 μl of lysis buffer was added to the cells and the mixture vortex vigorously for 15 seconds. All 400 μl of the lysate was transferred to well 1 of a Maxwell 16 LEV cartridge and 5 μl of DNase I solution added to well 4 of the cartridge. Elution tubes with 40–50 μl of nuclease-free water and LEV plungers were placed in the cartridge and then transferred to the Maxwell 16 Instrument the run started. RNA quality was measured using an Agilent TapeStation Instrument.

### qRT-PCR Assay to Monitor mRNA Target Knock-down Efficiency

Reverse transcription was performed using the iScript cDNA Synthsis Kit from BioRAD (Catalog number: 170–8891). Reactions were assembled containing 2μl of 5× iScript reaction mix, 0.5 μl of the iScript reverse transcriptase and 200 ng of the RNA in a total reaction volume of 10 μl. The reactions were incubated for 5 minutes at 25°C, 30 minutes at 42°C, 5 minutes at 85°C and then placed at 4°C. qPCR assays were assembled using Phusion High-Fidelity DNA Polymerase from NEB (Catalog number: M0530L) and SYBR Green from Invitrogen (Catalog number: S7563) and contained 4μl of 5X Phusion HF Buffer, 0.4 μl of 10 mM dNTPs, 1μl of 10 μM Forward Primer, 1 μl of 10 μM Reverse Primer, 1 μl of a 1:20 diulution of the cDNA reaction, 0.2 μl of Phusion DNA Polymerase, and 0.1 μl of 10,000X SYBR Green in a total volume of 20 μl. The reactions were then incubated at 98°C for 30 seconds followed by 35 cycles of 98°C for 10 seconds, 58–66°C (depending on primers used) for 15 seconds, and 72°C for 10 seconds in a BioRad qPCR machine. Depletion levels were calcluated using the 2 - ΔΔCt Method.

### Western Blot Assay to Monitor Protein Target Knockdown Efficiency

Cell pellets were resuspended in 100 μl Lysis buffer (50 mM Tris-HCl, pH 7.4; 100 mM NaCl; 1% NP-40; 0.1% SDS; 0.5% sodium deoxycholate) with protease inhibitor (Roche cocktail), vortexed vigorously, and incubated on ice at least 30 min. The lysate was spun at 18,000g for 20 min at 4°C. The supernatant was collected in a new tube on ice and the protein concentration determined using the Pierce BCA kit. 30–60μg of protein from each sample was diluted into 4X sample buffer and 10X reducing agent (Invitrogen NuPage reagents) and heated at 70°C for 10 min. The samples were loaded onto 4–12% Bis–Tris gels for proteins 10–220 kDa, or 8% Tris-glycine gels for proteins over 220 kDa along with 3μl of Licor Odyssey prestained molecular weight marker (928–4000). The gels were run at 200 V for about an hour, until the dye just runs off the gel (less for small molecular weight proteins) in 1X MOPS running buffer with 500 μl of antioxidant to the inner chamber of the buffer tank. The gels were transfered to PVDF for 30 min in transfer buffer, with methanol and antioxidant, using a BioRad Semi–dry transfer apparatus. The membranes were then blocked by incubation in 5–10 mls of Licor Blocking buffer for 1 hour at room temperature. The membranes were then incubated with the RNA binding protein primary antibody (0.2 μg/ml) and the loading control primary antibody (mfg recommended dilution) diluted in Licor block with 0.1% Tween 20 on a rocker at 4°C overnight. The next day, the membranes were washed 4 times for 5 min each in TBST. The membranes were then incubated with the appropriate secondary antibodies (Rockland Fluorescent TrueBlot anti-rabbit IgG IRDye800 (Catalog number 18–3216–32) for the RBP and Licor IRDye680 secondary antibody for the loading control) that were diluted according to the manufacturers instructions in Licor blocking buffer with 0.1% Tween 20 and 0.01% SDS on a rocker for 30–60 min, at room temperature. The membranes were then washed 4 times for 5 min each in TBST and rinsed once in TBS(noT) and scaned on a Licor Odyssey instrument.

